# Population genomics of a threespine stickleback tapeworm in Vancouver Island

**DOI:** 10.1101/2022.05.15.491937

**Authors:** Kum C. Shim, Jesse N. Weber, Catherine A. Hernandez, Daniel I. Bolnick

## Abstract

We surveyed the genetic structuring of 12 *Schistocephalus solidus* tapeworm populations from Vancouver Island (BC, Canada) using ddRAD sequencing and compared it to that of their threespine stickleback fish hosts. There were small but mostly significant genetic differences among the tapeworm populations. PCA results separated the populations by watershed (from the lakes where the tapeworms were collected), but we could not determine if the genetic structures seen were due to discrete models (i.e. watershed) or to a continuous model (i.e. isolation by distance). However, the tapeworm genetic differences were significantly smaller (P < 0.001) than those of the fish, which indicates that the parasite disperses more readily than their fish hosts.

## Introduction

Parasites as an important driving force on the evolution of their hosts is undisputed; for example, they can affect the rapid evolution of their hosts behavior, physiology, and immunology (Fumagalli et al. 2011, Ebert and Fields 2020, Weber et al. 2021, Weber et al. 2017). Moreover, parasites’ interactions with their hosts are thought to have played major roles in the maintenance of sexual reproduction and mating systems (Jaenike 1978, Hamilton 1980, Vergara et al. 20014), evolution of ploidy levels (Nuismer and Otto 2004), and mutational rates (M’Gonigle et al. 2009).

Given the importance of parasites in shaping physiology, ecology, and evolution, it is surprising that these organisms are understudied when compared to their free-living counterparts (Sprehn et al. 2015, Nuismer 2017a and 2017b). For instance, the population genomics of parasites and factors governing their dispersal and gene flow are lesser known relative to their hosts (Sprehn et al. 2015 and references therein). This is more palpable for parasites with complex life cycles infecting very disparate hosts during their different life stages. These host can be a mix of sedentary (less-dispersing) and very mobile organisms (Schmid-Hempel, P. 2011). For these complex life cycle parasites, it is thought that their more mobile host will determine their dispersal and genetic structure. This forms the basis of the vagile host hypothesis, which states that parasites with more mobile hosts will have less genetic structure than those with less mobile hosts, and in parasites with complex life cycles, the most vagile host will determent the genetic structure of the parasite (Nadler 1995, Criscione and Blouin 2004, Prugnolle et al. 2005). This means that parasites with only one host in their life cycle will have genetic structures similar to those of their hosts, while parasites with complex life cycles will have their genetic structures most similar to their most mobile hosts.

This vagile host hypothesis has been tested with some mix results in one-host parasites. For example, the high similarity in genetic structuring of the pocket gopher and its lice *Geomydooecus actuosi* reveals the parasite’s dispersal patterns matches that of its host, but for the field mice *Apodemus sylvaticus* and its parasitic nematode *Amblyomma americanum*, the nematode has much higher genetic structuring than its mice host, revealing there was more factors at play in the dispersal of the nematode than its hosts’ dispersal (Hafner 1983, Nieberding 2004).

For parasites with complex life cycles, the vagile host hypothesis has been tested with degrees of success. In the few studies on genetic structure and dispersal of complex life cycle parasites involving mainly trematode worm parasites, which parasitize a combination of sedentary (snails) and mobile vertebrate hosts (Schmid-Hempel 2011), their dispersal and genetic structuring have been mainly influenced by their more mobile hosts (Davies 1999, Jarne and Theron 2001, Criscione and Blouin 2004). However, a study on the tapeworm *Schistocephalus solidus* in Alaska, Sprehn et al. (2015) maintained that the movement of the vagile host had a lesser influence on this parasite’s genetic structure and dispersal due to the significant genetic differences among the tapeworm populations even among those in relative close proximity. However, their work was based on a short (759bp) mitochondrial sequence and eight microsatellite loci, which might have affected the power of their analyses. To validate the work of Sprehn et al. (2015) and to further test the robustness of the vagile hypothesis, we conducted another study using the tapeworm *S. solidus* tapeworm collected from a wide range of locations in Vancouver Island and used a larger sample of markers from across the tapeworm’s genome for a more comprehensive genetic structure survey.

The tapeworm *S. solidus* has a complex life cycle, infecting copepods in its first larval stage. Threespine stickleback fish is the obligate second intermediate host, which is infected by the tapeworm after consuming infected copepods (Barber 2013, Barber and Scharsack 2009, Dubinina 1980). The tapeworm final hosts are fish-eating birds such as loons. The tapeworm reproduces in the birds’ intestines, producing thousands of eggs, which are disseminated with the birds’ feces (Barber 2013, Barber and Scharsack 2009, Dubinina 1980). Thus, the tapeworm has a combination of mobile (birds) and less mobile (copepods and sticklebacks) hosts.

We hypothesize that the tapeworm should display small to no genetic structuring in Vancouver Island due to it having a highly mobile bird host in accordance with the vagile host hypothesis. To test this hypothesis, we first collected tapeworms from 12 lakes separated by various distances in Vancouver Island and investigated the degree of genetic structuring and differentiation among their populations. Then, we compared the helminths genetic structuring to that of sticklebacks, one of their lesser mobile hosts, to see congruency between their patterns. If the parasite population genetic differentiation is significantly lower than that of their fish hosts, then the tapeworm’s mobile bird host is likely to be the main drive on the dispersal and genetic differentiation in the parasite.

We also tested if there was yearly turnover of tapeworm populations in the lakes in Vancouver Island using tapeworms collected in the same three lakes (Boot, Gosling, and Merrill lakes) but in different years (2012 and 2016). Sprehn et al. (2015) also found evidence of temporal variation in the genetic differentiation of the Alaskan tapeworm populations. They hypothesized that small effective population sizes and genetic drift from stochastic infection, reproduction, and dispersal events by their bird hosts might be the cause. However, they arrived at this conclusion by using different tapeworm populations from different years and analyzed their data on a hierarchical Analysis of Molecular Variance (AMOVA) using year as one of their groupings. We believe it is more appropriate to compare the genetic makeup of the same populations from different time periods to test if indeed there is temporal variation in the genetic differentiation of the tapeworms.

Results from this work indicated that there were small but significant genetic differences between the tapeworm populations in Vancouver Island, especially for populations that were highly apart (by more than 100Km). But these genetic differences were significantly lower than that of their stickleback hosts, indicating the parasite’s higher dispersal rate probably due to their mobile bird hosts. Lastly, there were no significant genetic differences between the tapeworm populations collected in 2012 to those collected in 2016, indicating a lack of temporal genetic differences at least in the three lakes (Boot, Gosling, and Merrill) populations.

## Materials and Methods

### Tapeworm collecting

The *S. solidus* tapeworms were collected in June and July 2016 and 2017. For more details on the locations and sample sizes, see table 1 and figure 1. The threespine stickleback fish were collected using un-baited minnow traps left submerged overnight in shallow water (< 3m) along each lake’s shorelines (Scientific Fish Collection Permit No. NA16-230545 [2016] and MRNA16-263390 [2017], Ministry of Forest, Lands and Natural Resource Operations, BC, Canada). A random sample of captured fish were euthanized in MS-222 and dissected on-site (methods approved by The University of Texas Institutional Animal Care and Use Committee as part of AUP-2010-00024). The size (i.e. weight and length), sex, and tapeworm infection status of each fish were recorded during dissection. The tapeworms were stored in 1.5mL centrifuge tubes (one per fish) and preserved in 100% ethanol for later DNA extraction.

**Table 1:** *Schistocephalus solidus* tapeworm sample locations, collection dates, sample sizes, infection levels in threespine stickleback fish, and watersheds.

| Lake | Latitude | Longitude | Collection date | Fish N<sup>1</sup> | Tapeworm N<sup>2</sup> | Infection rate (%)<sup>3</sup> | watershed |
| --- | --- | --- | --- | --- | --- | --- | --- |
| Frederick | 48.858001 | -125.023058 | 6/29/2016 | 30 | 22 | 80.0 | Alberni Inlet |
| Pachena | 48.837661 | -125.033891 | 7/2/2016 | 7 | 25 | 71.4 | Alberni Inlet |
| Blackwater | 50.165982 | -125.593771 | 6/1/2017 | 91 | 17 | 8.1 | Amor Lake |
| Farewell | 50.199494 | -125.586098 | 6/25/2016 | 46 | 24 | 16.7 | Amor Lake |
| Boot (2012) | 50.055025 | -125.526444 | 6/30/2012 | 20 | 24 | 95.0 | Campbell River |
| Boot | 50.055025 | -125.526444 | 6/24/2016 | 31 | 25 | 87.1 | Campbell River |
| Echo | 49.987652 | -125.411327 | 6/19/2016 | 41 | 25 | 43.9 | Campbell River |
| Gosling (2012) | 50.045922 | -125.5014 | 6/28/2012 | 16 | 14 | 78.6 | Campbell River |
| Gosling | 50.045922 | -125.5014 | 6/24/2016 | 39 | 24 | 41.9 | Campbell River |
| Higgins | 50.070486 | -125.508545 | 6/7/2017 | 83 | 22 | 21.9 | Campbell River |
| Lawson | 50.035596 | -125.578308 | 6/24/2016 | 41 | 24 | 29.0 | Campbell River |
| Merrill (2012) | 50.060494 | -125.550094 | 7/1/2012 | 27 | 23 | 94.7 | Campbell River |
| Merrill | 50.060494 | -125.550094 | 6/23/2016 | 44 | 24 | 21.2 | Campbell River |
| Comida | 50.15056 | -125.52813 | 6/3/2017 | 195 | 23 | 4.4 | Mohun Lake |
| Roselle | 50.524178 | -126.989632 | 6/20/2016 | 61 | 25 | 31.1 | Nimkish Lake |

<sup>1</sup> Number of threespine stickleback fish collected for this work.
<sup>2</sup> Number of *Schistocephalus solidus* tapeworms used per lake for this work from a sample of the previous fish N column.
<sup>3</sup> Tapeworm infection rate in the threespine stickleback fish from each lake.

**Figure 1:**
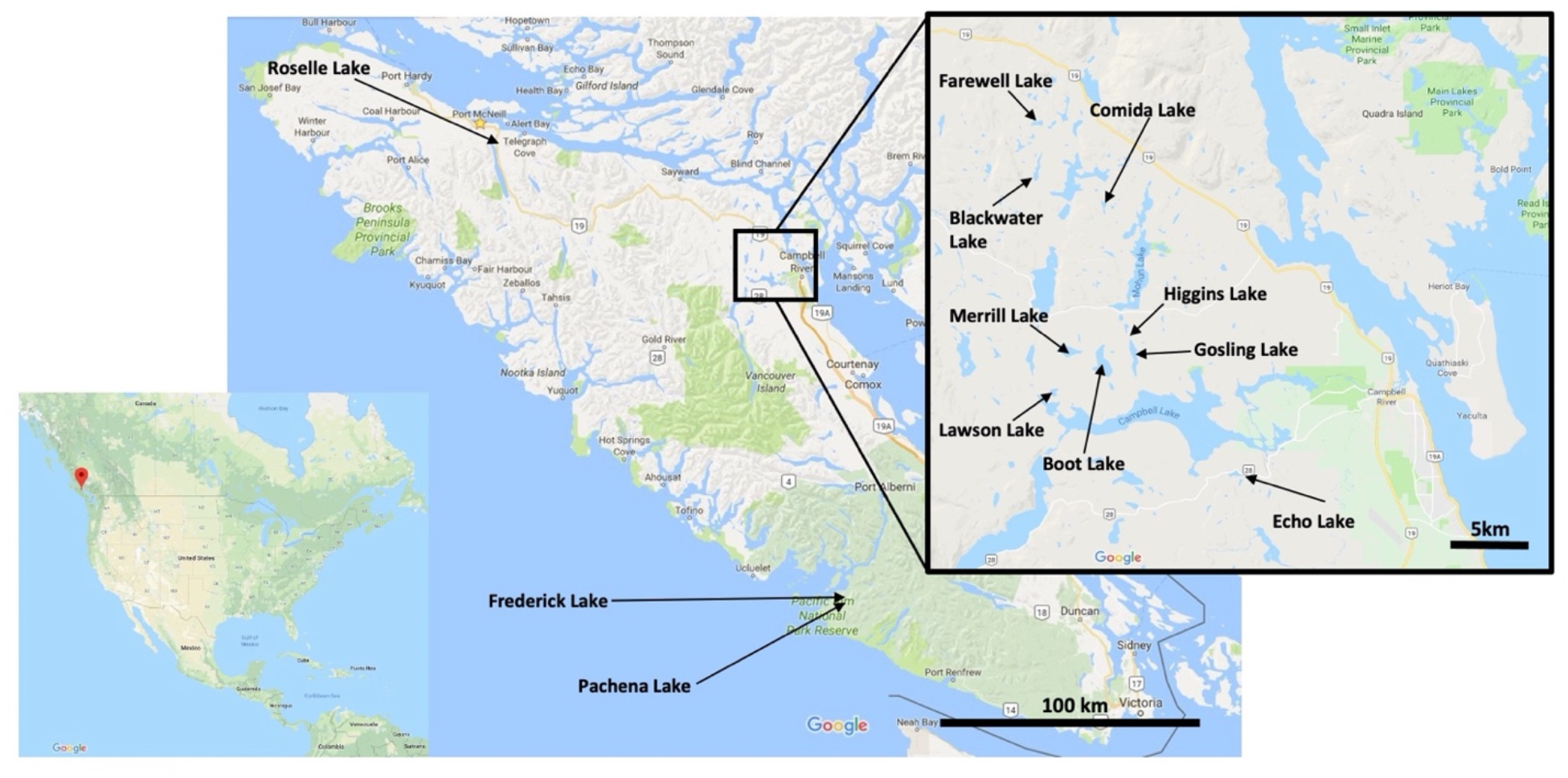
Lake locations in Vancouver Island.

Tapeworms from three populations (Boot, Gosling, Merrill lakes) collected in 2012 by Catherine Rodriguez (a former undergraduate research assistant) was added to this work to see if there were genetic differences between different years in the same populations. See table 1 for collection and sample size details.

### DNA extraction and ddRAD sequencing

The DNA for each tapeworm was extracted back in our laboratory in Texas using the Wizard® SV 96 Genomic DNA Purification Kit and System by Promega (Madison, WI). The DNA concentration was quantified using Quan-iT PicoGreen dsDNA Assay Kits and dsDNA Reagents by Thermofisher (Waltham, MA) and a 96-well microplate reader (Tecan Infinite M200 Pro, Mannedorf, Switzerland). After quantification, ∽15 μL of DNA solution per sample (containing ∽240ng of DNA) were used for ddRAD processing.

Double digest Restriction-site Associated DNA (ddRAD) is a variation in RAD sequencing methodology whereby restriction two endonuclease enzymes are used to cut an organisms’ whole genome into consistent fragments. This is a cheap and effective methodology for sampling organisms’ genome for SNP discovery and genotyping (Peterson et al. 2012).

We followed the protocols from Peterson at al. (2012) and Stuart et al. (2017) for extracting and processing paired end ddRAD fragments, but we used SpHI and EcoRI restriction endonucleases instead for this work. From simulations using simRAD (an R [R Core team] package to simulate the number of loci expected in RAD sequencing using predicted restriction endonucleases [Lepais and Weir 2014]), we expected to obtain ∽9,000 loci for 300-400bp ddRAD fragments for the tapeworm, which has a sequenced genome of ∽540Mbp in size (WormBase in Howe et al. 2017).

The filtered 350-400bp ddRAD fragments (420-470bp if including the barcode tags for each fragment) were sent to the College of Natural Sciences Genome Sequencing and Analysis Facility (GSAF, Austin, TX) for sequencing. The sequencer used was an Illumina HiSeq 4000, using 2 × 150bp read lengths for an average depth coverage of ∽723 reads per site.

### Bioinformatics

The raw sequenced files were trimmed from the Illumina adaptors and demultiplexed (i.e. sorting and assigning names to the sequenced reads) using the *process_radtags* function from the program Stacks v2.0 (Catchen et al. 2013).

We used the dDocent bioinformatic pipeline (Puritz et al. 2020) for filtering and SNP-calling on the demultiplexed ddRAD sequences, obtaining a final compiled and filtered Variant Call Format (VCF) file for my sequences. Ten individual tapeworms were removed during VCF filtering due to >50% missing data. We used the *S. solidus* tapeworm reference genome from the Aubin-Horth group, Université Laval (Berger et al., 2021) to map the ddRAD sequences.

We also modified the dDocent script and reran the pipeline with the demultiplexed ddRAD sequences to obtain a Genomic VCF (GVCF) file, which contains variant and in-variant loci for calculating the tapeworm heterozygosity and nucleotide diversity (π). Using VCF file, which only contains the variant loci would bias the heterozygosity and π results upward (i.e. bigger than real values), hence a GVCF file is recommended for these calculations (Korunes and Samuk 2021, Dan Bolnick in personal communications). We used the *populations* function from the program Stacks v2.6 (Catchen et al. 2013) to calculate the population summary of genetic diversity statistics, including heterozygosity and π using the GVCF file.

The VCF file was transformed into an Arlequin v3.5.2.2 program file (i.e. arp file; Excoffier and Lischer 2010) using the PGDSpider v2.1.1.5 program (Lischer and Excoffier 2012). We used Arlequin to find outlier loci (using the Hierarchical Island Model and 50,000 coalescent simulations as settings), which we removed from the VCF file using the R package vcfR v1.12.0 (Knaus and Grünwald 2017) before downstream data analyses. After removing the outlier loci, we also used Arlequin to calculate pairwise F_ST_ (using 50,000 permutations for significant test) among the populations and to perform an Analysis of Molecular Variance (AMOVA; using 16,000 permutations for significant test for the F statistics) for the 15 tapeworm populations. We used the R program Ape v5.5 (Paradis et al. 2004) to construct a neighbor joining tree for the tapeworm populations using the F_ST_ values.

We used the R program Adegenet v2.1.5 (Jombart 2008) to perform the Principal Components Analysis (PCA), Discriminant Analysis of Principal Components (DAPC) to cluster the tapeworm populations, and to test Isolation by Distance (IBD) between the populations using the Mantel Test.

We used the R program ConStruct v1.0.4 (Bradburd et al. 2018) using the allele frequencies from SNPs obtained from Arlequin. We used ConStruct to infer if the tapeworm population structures follow continuous (i.e. IBD) or discrete models. This program compares the predictive accuracy of both models to determine which describes best the given genetic data. We used the following settings for this program: 1 to 8 layers (K), eight cross-validation replicates to be run, and 10,000 iterations per Markov Chain Monte Carlo (MCMC) run. Only bi-allelic SNPs were used for this analysis as per the manual.

Lastly, we compared in R the stickleback fish and tapeworm F_ST_ from the same lakes but collected in different years (2014 for the fish and 2016-17 for the tapeworms). The stickleback F_ST_ data is from Stuart et al. (2017). The host and parasite’s similarity in genetic differentiation (i.e. F_ST_) would indicate a close host-parasite association at least on dispersal rates.

## Results

The ddRAD sequencing of 341 tapeworms resulted in ∽20,000,000 read pairs. After SNP-calling and quality filtering using dDocent, 8,234 SNPs out of possible 1,386,954 were retained for downstream analyses. This number of retained SNPs was close to the predicted 9,000 by the simRAD R package. We found at least 21 outlier SNPs after outlier loci test in Arlequin. These are shown in supplementary table 1.

The original GVCF file used to estimate genetic diversity (observed and estimated heterozygosity [H_O_ and H_E_], π, and inbreeding coefficient [F_IS_]) had 2,345,453 variant and invariant loci, but the Stacks v2.6 program automatically filtered and kept only 933,198 loci (507,063 of which were variant according to the program) to estimate the genetic diversity statistics. All the tapeworm populations had similarly consistent genetic diversity indices (H_O_ ranged from 0.011 to 0.016, H_E_ from 0.031 to 0.061, and π ranged from 0.036 to 0.067 [table 2]), except for F_IS_, which ranged from 0.054 to 0.137. All the populations also had consistent low percent polymorphic loci, which ranged from 12% to 23% (table 2).

**Table 2:** Population summary of genetic diversity statistics

| Lakes | All Sites<sup>1</sup> | Variant Sites<sup>2</sup> | % Polymorphic Loci<sup>3</sup> | H<sub>o</sub><sup>4</sup> | H<sub>E</sub><sup>5</sup> | Pi<sup>6</sup> | F<sub>IS</sub><sup>7</sup> |
| --- | --- | --- | --- | --- | --- | --- | --- |
| Blackwater | 609277 | 379461 | 11.866 | 0.012 | 0.031 | 0.036 | 0.057 |
| Boot (2012) | 635884 | 397267 | 14.745 | 0.013 | 0.036 | 0.040 | 0.070 |
| Boot | 667681 | 423547 | 16.407 | 0.014 | 0.041 | 0.046 | 0.084 |
| Comida | 637155 | 402586 | 14.572 | 0.011 | 0.037 | 0.042 | 0.078 |
| Echo | 744611 | 471048 | 22.566 | 0.016 | 0.061 | 0.067 | 0.137 |
| Farewell | 645909 | 409402 | 13.981 | 0.011 | 0.037 | 0.041 | 0.077 |
| Frederick | 634225 | 399025 | 14.705 | 0.012 | 0.037 | 0.041 | 0.077 |
| Gosling (2012) | 619618 | 387182 | 11.954 | 0.014 | 0.033 | 0.038 | 0.054 |
| Gosling | 659963 | 416573 | 14.686 | 0.012 | 0.038 | 0.043 | 0.078 |
| Higgins | 719203 | 460052 | 18.274 | 0.015 | 0.052 | 0.058 | 0.105 |
| Lawson | 615403 | 385172 | 13.553 | 0.011 | 0.034 | 0.037 | 0.071 |
| Merrill (2012) | 636841 | 395999 | 14.607 | 0.012 | 0.035 | 0.039 | 0.071 |
| Merrill | 630996 | 389848 | 14.494 | 0.012 | 0.035 | 0.039 | 0.072 |
| Pachena | 719171 | 458350 | 19.116 | 0.015 | 0.051 | 0.058 | 0.108 |
| Roselle | 612090 | 375792 | 13.254 | 0.012 | 0.031 | 0.035 | 0.060 |

<sup>1</sup>All sites include variant and non-variant sites from the GVCF (genomic VCF) file used.
<sup>2</sup>Variant sites indicate the number of SNPs found by Stacks v.2.6 from the unfiltered gvcf.
<sup>3</sup>Proportion of SNPs in each population/lake.
<sup>4</sup>H<sub>o</sub> is observed heterozygosity.
<sup>5</sup>H<sub>E</sub> is expected heterozygosity
<sup>6</sup>Pi is nucleotide diversity
<sup>7</sup>F<sub>IS</sub> is the inbreeding coefficient of individuals relative to the subpopulation.

The pairwise F_ST_ between the lakes ranged from −0.0004 (for Gosling Lake tapeworms collected in 2016 and 2012) to 0.096 (for Farewell and Roselle lakes [table 3]). Most of the F_ST_ were significantly different (P < 0.005) except for the comparisons of tapeworm populations collected in the same lake but in different years (i.e. Boot 2016 vs Boot 2012, Gosling 2016 vs Gosling 2012, and Merrill 2016 vs Merrill 2012) and between lakes that were in close proximity (i.e. Boot, Gosling, Higgins, and Merrill). Overall, lakes that were in farther apart had the highest F_ST_; the lake distances unsurprisingly also coincide with that of their watersheds. The neighbor joining tree based on the F_ST_ values illustrates this result: lakes that were in close geographical proximity (or belonging to the same watershed) were sister groups (figure 2). The hierarchical AMOVA results indicated a strong and significant difference among the five watersheds (F_CT_ = 0.052, P<0.0001, table 4).

**Table 3:** Pairwise F_ST_ for the 15 tapeworm population lake comparisons.

|  | Blackwater | Boot (2012) | Boot | Comida | Echo | Farewell | Frederick | Gosling (2012) | Gosling | Higgins | Lawson | Merrill (2012) | Merrill | Pachena | Roselle |
| --- | --- | --- | --- | --- | --- | --- | --- | --- | --- | --- | --- | --- | --- | --- | --- |
| Blackwater | — | — | — | — | — | — | — | — | — | — | — | — | — | — | — |
| Boot (2012) | **0.055** |  |  |  |  |  |  |  |  |  |  |  |  |  | — |
| Boot | **0.055** | 0.002 |  |  |  |  |  |  |  |  |  |  |  |  | — |
| Comida | **0.039** | **0.018** | **0.017** |  |  |  |  |  |  |  |  |  |  |  | — |
| Echo | **0.052** | **0.007** | **0.006** | **0.018** |  |  |  |  |  |  |  |  |  |  | — |
| Farewell | **0.008** | **0.065** | **0.065** | **0.049** | **0.062** |  |  |  |  |  |  |  |  |  | — |
| Frederick | **0.061** | **0.059** | **0.057** | **0.039** | **0.052** | **0.068** |  |  |  |  |  |  |  |  | — |
| Gosling (2012) | **0.056** | 0.001 | 0.001 | **0.016** | **0.006** | **0.067** | **0.058** |  |  |  |  |  |  |  | — |
| Gosling | **0.058** | 0.002 | 0.001 | **0.017** | **0.007** | **0.067** | **0.059** | -4E-4 |  |  |  |  |  |  | — |
| Higgins | **0.049** | **0.004** | **0.003**<sup>3</sup> | **0.014** | **0.007** | **0.059** | **0.053** | 0.001 | 0.002 |  |  |  |  |  | — |
| Lawson | **0.058** | **0.004**<sup>1</sup> | **0.005** | **0.021** | **0.010** | **0.068** | **0.062** | **0.005**<sup>4</sup> | **0.005** | **0.006** |  |  |  |  | — |
| Merrill (2012) | **0.056** | 0.001 | 0.001 | **0.017** | **0.006** | **0.066** | **0.058** | -0.001 | 0.001 | 0.001 | **0.005** |  |  |  | — |
| Merrill | **0.058** | **0.003**<sup>2</sup> | 0.001 | **0.019** | **0.007** | **0.067** | **0.060** | 0.001 | 0.001 | **0.003** | **0.004** | 0.001 |  |  | — |
| Pachena | **0.058** | **0.057** | **0.057** | **0.040** | **0.052** | **0.067** | **0.005** | **0.058** | **0.059** | **0.052** | **0.061** | **0.057** | **0.059** |  | — |
| Roselle | **0.090** | **0.084** | **0.080** | **0.068** | **0.077** | **0.096** | **0.075** | **0.085** | **0.084** | **0.077** | **0.086** | **0.083** | **0.083** | **0.075** | — |

Note: Results below are from Arlequin 3.5.2.2 (Population pairwise differences ( $F_{ST}$ ) based on Nei & Li (1979), gamma = 0.01, and 10,100 permutations for the significance level test).
P values for all significant differences (those in bold) are $P < 0.005$ except the following:
<sup>1</sup>Boot (2012) and Lawson: P value = 0.00554+/-0.0007
<sup>2</sup>Boot (2012) and Merrill: P value = 0.01188+/-0.0011
<sup>3</sup>Boot and Higgins: P value = 0.01+/-0.0011
<sup>4</sup>Gosling (2012) and Lawson: P value = 0.03148+/-0.0016

**Table 4:** Analysis of Molecular Variance (AMOVA) results for the tapeworm populations from five watersheds and among populations.

| Source of Variation | Sum of Squares | Variance Components | Percent Variation | Fixation indices |
| --- | --- | --- | --- | --- |
| Among the 5 watersheds | 29366.04 | 59.53 | 5.21 | $F_{CT} = \mathbf{0.052}$ |
| Among populations within the 5 watersheds | 12940.41 | 4.81 | 0.42 | $F_{SC} = \mathbf{0.004}$ |
| Among the 15 populations | 71.8578.49 | 1077.33 | 94.36 | $F_{ST} = \mathbf{0.056}$ |

Note: All *F-statistics* significant ( $P < 0.0001$ )

**Figure 2:**
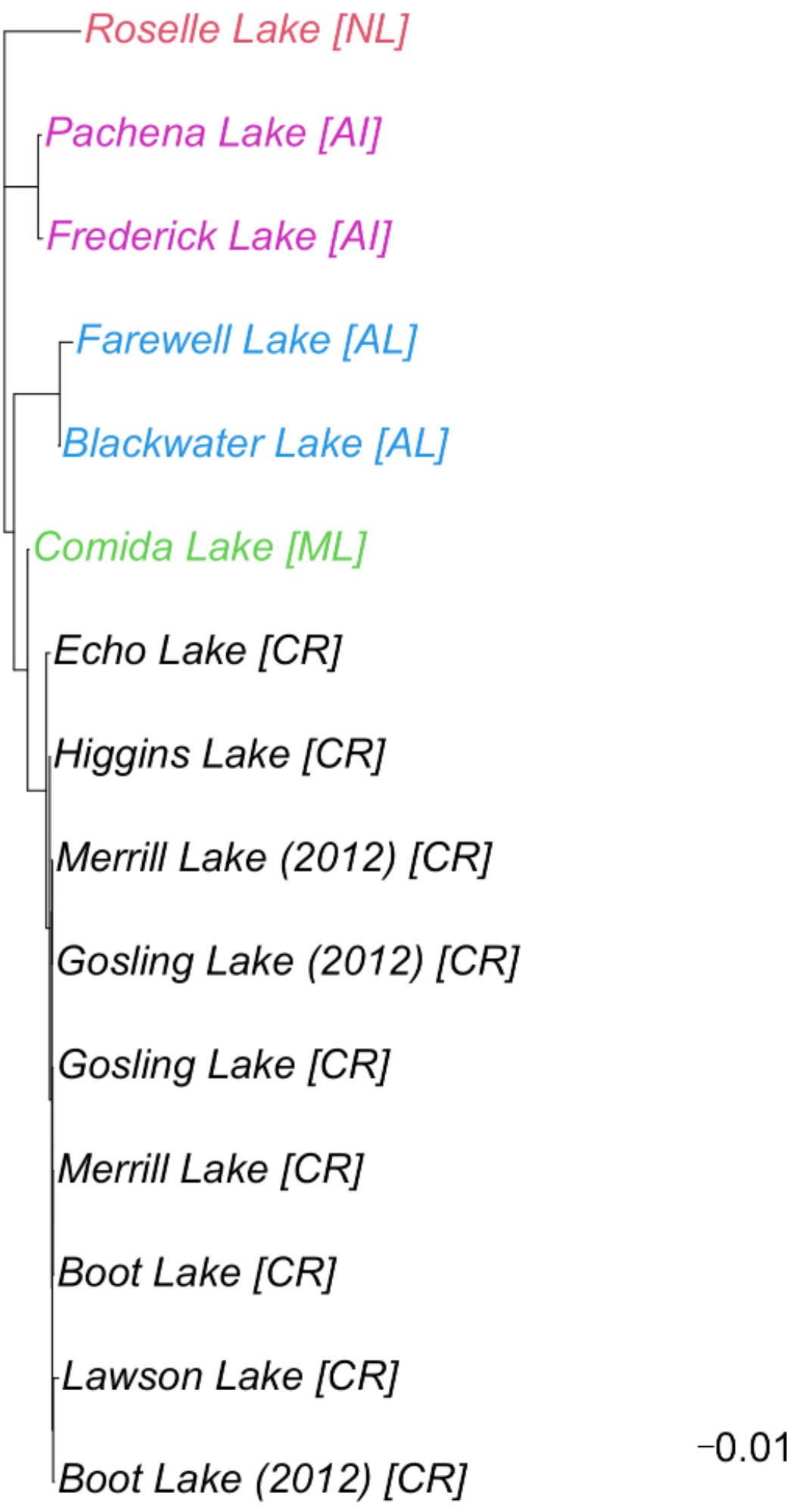
Neighbor Joining tree (from pairwise F_ST_ values) for the tapeworm populations from each lake. Purple colored lakes are from Alberni Inlet Watershed (AI), red lake is from Nimpkish Lake Watershed (NL), blue for Amor Lake Watershed (AL), green for Mohun Lake Watershed (ML), and black for Campbell River Watershed (CR). The initials of the watersheds are also enclosed in brackets next each lake’s names in the tree.

The PCA indicated five discrete groupings, which coincided with watershed (and/or geographical proximity [figure 3]). However, the DAPC analyses estimated K = 3 as the best number of clusters to describe the structuring of the tapeworm populations, but with K = 2 to 5 close behind (supplementary figure 1). We provided plots representing K= 2 to 5 (figure 4), with the K = 5 providing the most similar results to PCA (figure 3) as in having each watershed assigned to its own cluster (figure 4). For K = 3, all populations from Nimpkish Lake and Alberni Inlet watersheds (tapeworm populations from Roselle, Frederick, and Pachena lakes) clustered into one group, Campbell River Watershed lakes clustered into another group, while lakes from Amor Lake and Mohun Lake watersheds (Blackwater, Farewell, and Comida lakes) clustered into a group with considerable probability of also belonging to the Campbell River Watershed group (figure 4). See supplementary table 2 for the alpha scores and the suggested number of PCs used for each K value in the DAPC analyses.

**Figure 3:**
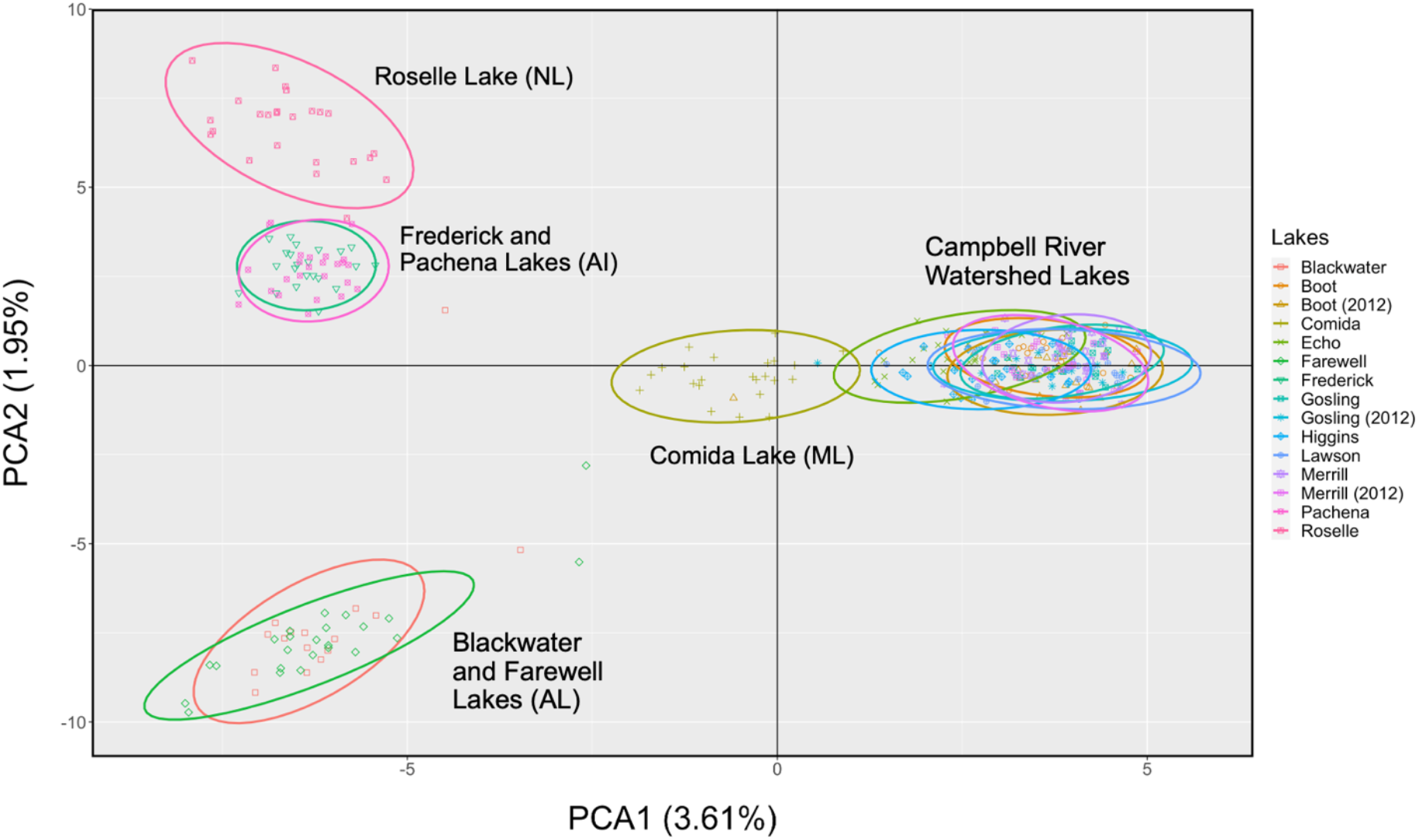
PCA plot using the first two principal components based on all SNP loci. The initials of watershed names are in parenthesis next to lake names in graph, and they are: AI: Alberni Inlet Watershed, AL: Amor Lake Watershed, ML: Mohun Lake Watershed, NL: Nimpkish Lake Watershed.

**Figure 4:**
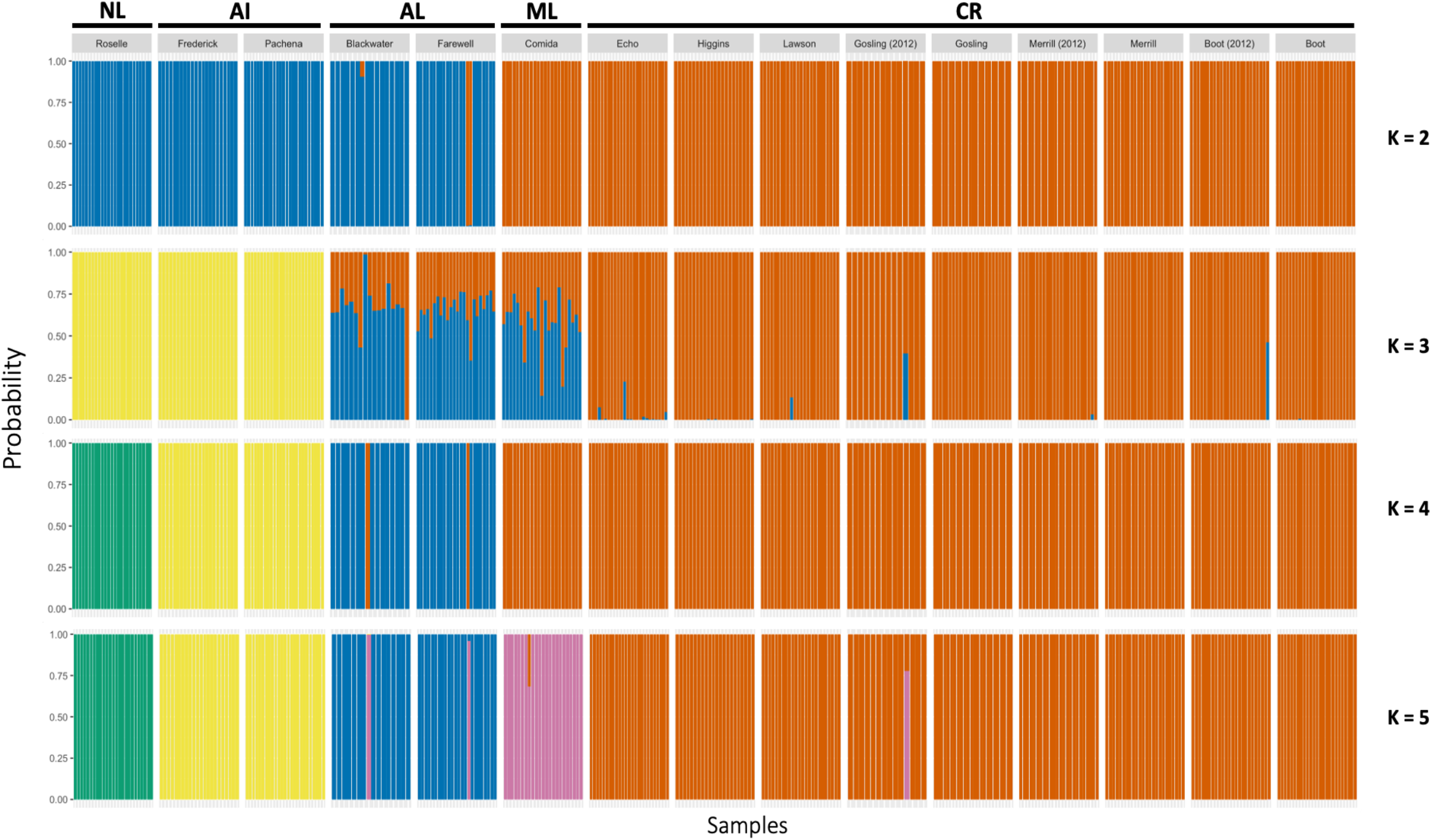
Individual clustering based on DAPC results from K = 2 to 5. The initials of watershed names are above each lake’s names, and they are: NL (Nimpkish Lake Watershed), AI (Alberni Inlet Watershed), AL (Amor Lake Watershed), ML (Mohun Lake Watershed), CR (Campbell River Watershed).

The Mantel Test strongly indicated Isolation by Distance (IBD) as the cause of the genetic variation in the tapeworm populations (P = 0.002, figure 5). However, the watersheds that the tapeworm populations belong to are also separated by similar distances (see figure 1). Analysis with ConStruct indicated IBD had higher predicted accuracy for K = 1 to 2 than discrete clustering models (figure 6); however, there are no significant differences when K is three and above (figure 6), meaning it was unable to discern whether IBD or discrete models best described the data.

**Figure 5:**
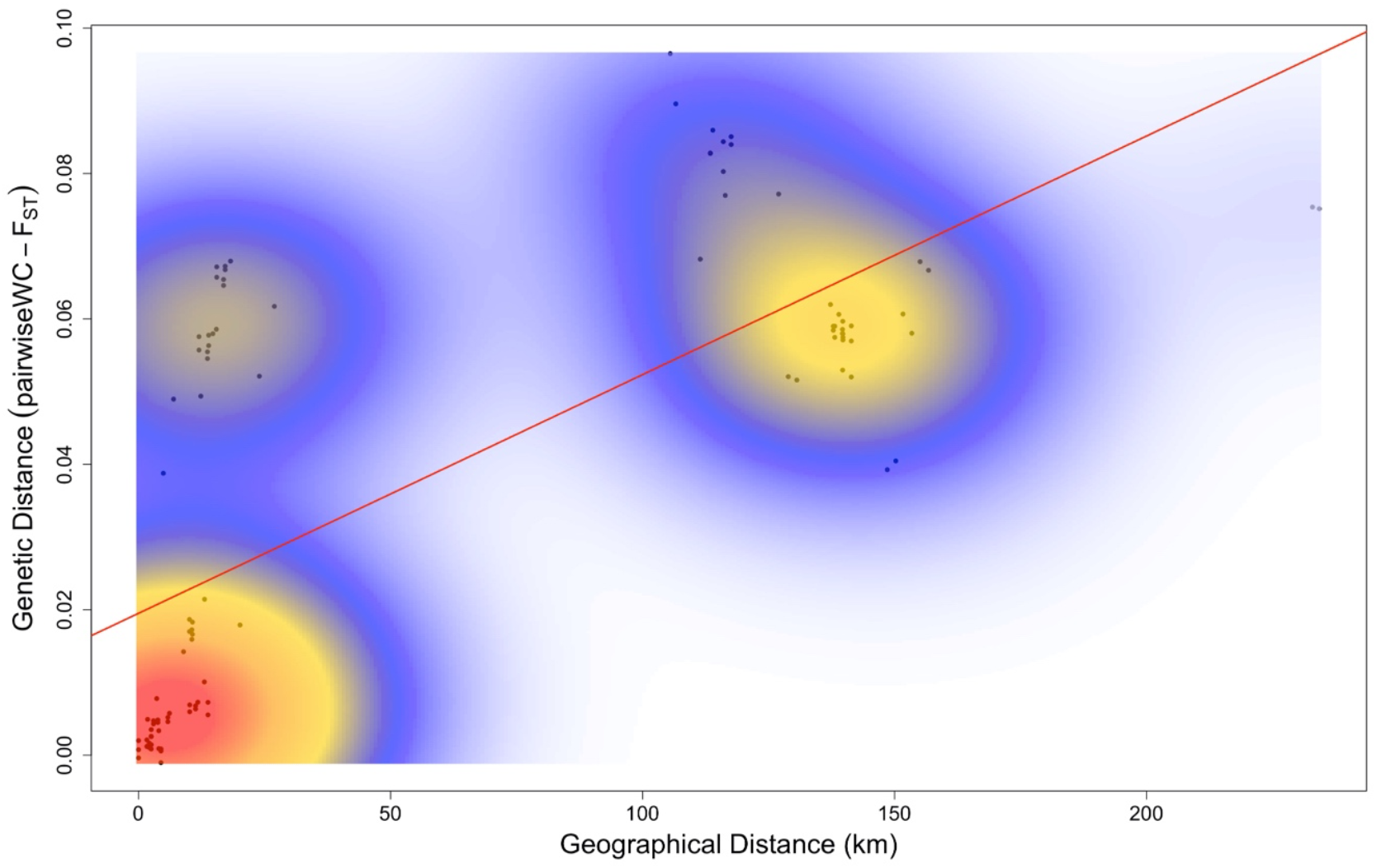
There was strong correlation between genetic distance and geographical distance indicating likely Isolation by Distance (IBD) as the main cause in the natural variation of the tapeworm populations. This is also indicated by the Mantel Test for Isolation by Distance (P = 0.002). However, these results are confounded by the watersheds that the tapeworm populations belong to since these are also separated by the same distances. The color gradient indicates density of points with red as the densest.

**Figure 6:**
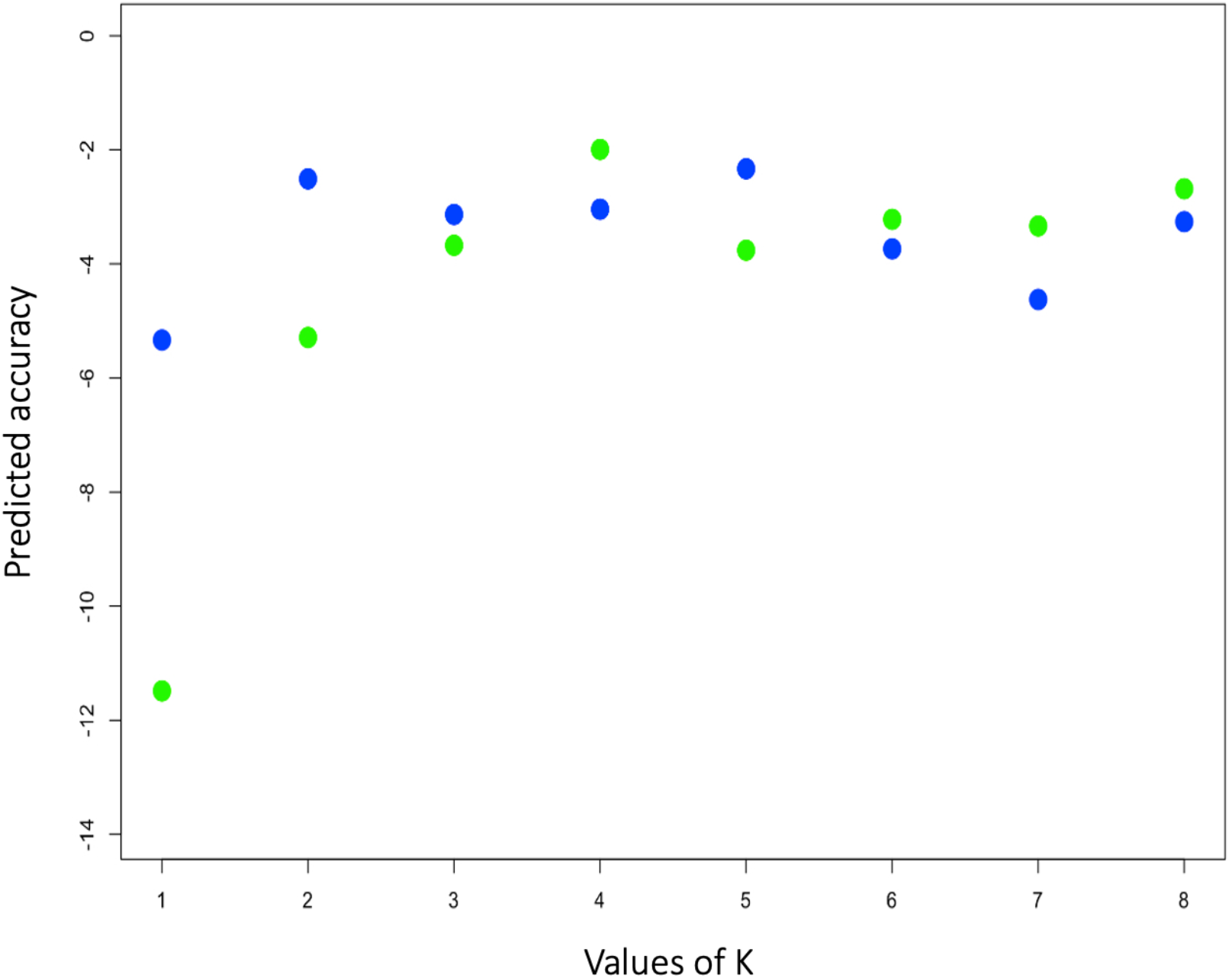
Result of analysis using ConStruct v1.0.4 indicated IBD model (blue dots) as the best predictor (i.e. closest to zero in the predicted accuracy scale) when K = 1 to 2, but it is not different to discrete clustering models (represented by green dots) for K >2.

The stickleback fish and tapeworm populations both follow positive correlation trends between distance among populations and increasing F_ST_ (figure 7). However, the trend was steeper in the fish populations due to them having significantly higher F_ST_ (P<0.0001). The relative slopes of the two regressions were also significantly different (P <0.0001; see supplementary material for details of the slope comparison).

**Figure 7:**
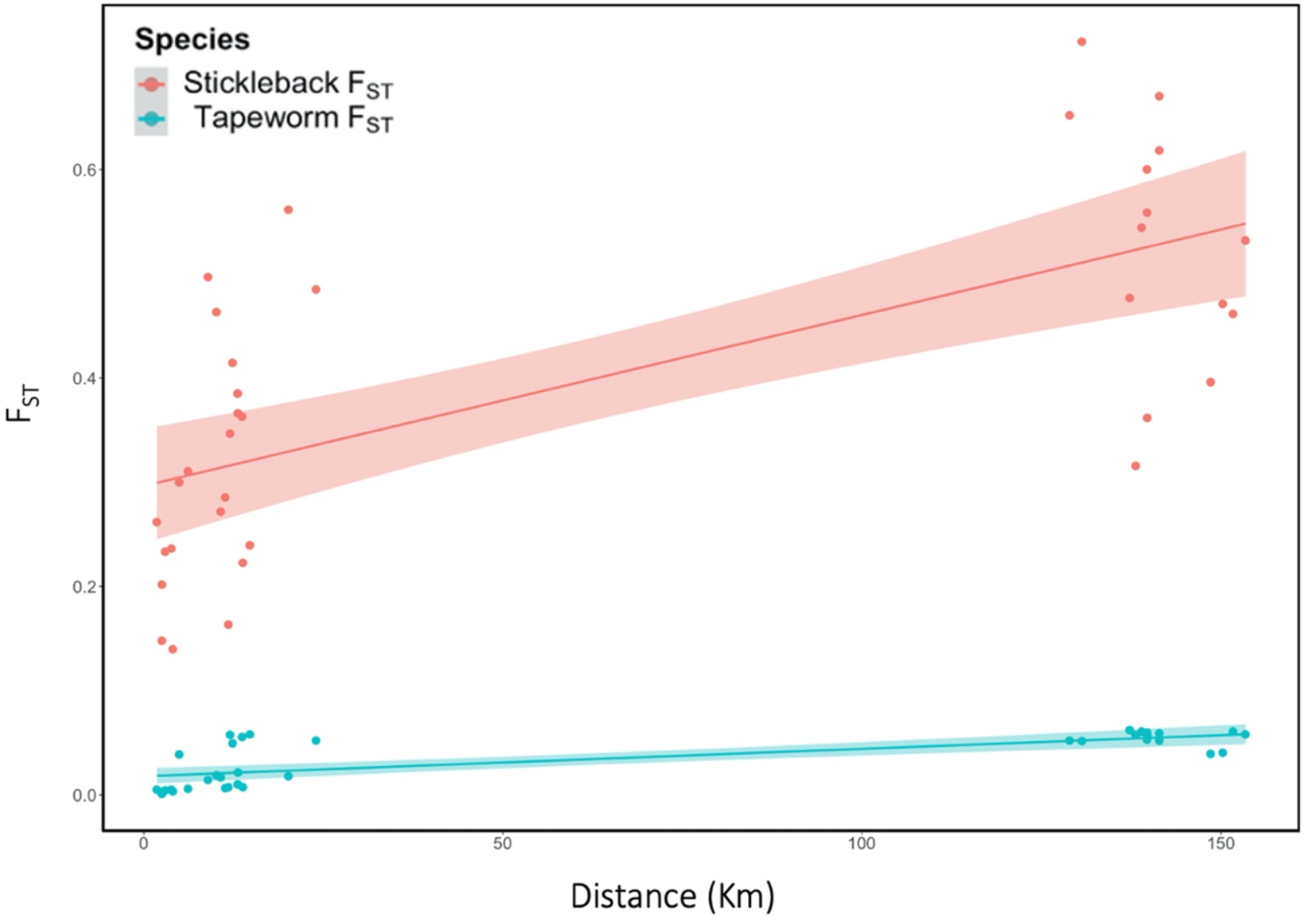
Threespine stickleback fish and *S. solidus* tapeworm populations both have positive correlations between increasing F_ST_ and increasing distance among the populations. However, the trend is steeper in the fish due to it having significantly higher genetic differences (F_ST_) among its populations than the tapeworm (P<0.0001).

## Discussion

For this project, we tried to elucidate the genetic structuring of the *S. solidus* tapeworm populations in Vancouver Island, compare this structuring to that of their stickleback host and test for the validity of the vagile host hypothesis, which states that the more mobile host should determine the genetic structuring of their parasites (Nadler 1995, Criscione and Blouin 2004, Prugnolle et al. 2005).

The genetic differences (measured by F_ST_) between most tapeworm populations in Vancouver Island was relatively small but significant except for a few populations from lakes that were in close proximity (i.e. Gosling, Higgins, Boot, and Merrill Lakes [table 3]); this indicates that there were relatively more gene flow between nearby tapeworm populations. Similarly, past work on the tapeworm populations in Alaska indicated that there was also significant genetic structuring of this parasite in that region (Sprehn et al. 2015). However, the overall F_ST_ among tapeworm populations was much higher in Sprehn et al. (2015), with some having over an order of magnitude higher (i.e. Sprehn et al. [2015] had a maximum F_ST_ = 0.142 vs the maximum F_ST_ = 0.096 for my work). Also, some closely distanced lakes in Sprehn et al. (2015) were also highly different (e.g., two lakes separated by ∽1km in Alaska had F_ST_ = 0.108 [Sprehn et al. 2015]), while lakes from my work in Vancouver Island that were 1-10km apart, had a range of FST = 0.001 to 0.007 (see table 3 and figure 1). If the results from Sprehn et al. (2015, using genetic data from a 759bp mitochondrial sequence and 8 microsatellite loci) are robust, then the genetic structuring of the tapeworm populations in Alaska are very different from those in Vancouver Island, probably due to different processes at play. Sprehn et al. (2015) hypothesized that perhaps discrete territoriality or migratory patterns of loons (which are the main bird host for the tapeworm in both Alaska and Vancouver Island [Sprehn et al. 2015, personal observations]) might account for these relatively high F_ST_ for nearby lakes in that region in Alaska.

There were also no genetic differences for the Vancouver Island tapeworm populations collected in different years (between 2012 and 2016 for Boot, Gosling, and Merrill lakes [table 3]), indicating no population turnover between the years. Again, this result is different from Sprahn et al. (2015), who claimed there were temporal differences in the genetic structuring of their tapeworm populations when using collecting year as a grouping in their AMOVA analyses. However, they did not collect the same populations in their various collecting years (but different populations in different years); thus, their results might be confounded by structuring between the populations instead of representing differences over the years in the same populations.

My results also mostly support the vagile host hypothesis in that complex life cycle parasites with sedentary and mobile hosts, will have their genetic structure mainly determined by their mobile host. The sticklebacks and tapeworms F_ST_ were both positively correlated with distance between lakes (figure 7), but the correlation was steeper in the fish, and their F_ST_ values were significantly higher (P<0.0001) than those of the parasite populations. This indicates the tapeworms had higher dispersal than their fish hosts, dispersal which is likely due to their final bird hosts. Stickleback populations in Vancouver Island lakes are genetically highly structured (Caldera and Bolnick 2008, Stuart et al. 2017, figure 7) and are not known to migrate much into other lakes. Even populations from streams are very different from those in lakes connected to said streams (Stuart et al. 2017, Izen et al. 2016, and personal observations). Although there is no work on the dispersal and genetic structure of copepods (the parasites’ first host) in Vancouver Island lakes, works with other species confined to specific bodies of water from elsewhere indicates that these crustaceans are also highly non-migratory (Burton and Feldman 1981, Young et al. 2013, Zhang et al. 2013). Loons are the primary fish-eating birds in Vancouver Island lakes in the summer (personal observations) when the tapeworms are at their prime bird-infective stage (Dubinina 1980, personal observations), and they are known to be highly mobile, having several territorial lakes for feeding during mating season and might roam to more lakes when not reproducing (Barr 1996, Piper et al. 1997). Thus, for the relatively low genetic differences found in the Vancouver Island tapeworms, the highly mobile bird hosts should be the main dispersal agent for the parasite, supporting the vagile hypothesis.

Even though the genetic differences between the tapeworm populations in Vancouver Island were small, most of them were still significantly different (table 3), so more processes besides dispersal via mobile bird host might be at play. Ecological factors like lake water chemistry or temperature might affect the survival of some tapeworm genotypes, causing some genetic structuring in the tapeworm populations. This is because the complex life cycle of the tapeworm involves deposition of its eggs in lakes for an unknown period of time and a brief (∽24hr) free living larval stage (Dubinia 1980) in the water. Another possibility is that copepods and stickleback hosts, which are genetically different between lakes (Stuart et al. 2017, Caldera and Bolnick 2008) might also select for certain tapeworm genotypes in different lakes, causing again genetic differentiation between the tapeworm populations. Thus, all these physical and biological factors might act together to produce a pattern of small but significant genetic differences in the tapeworm populations, especially for those that are moderately to highly geographically apart. Sprehn et al. (2015) came to similar thoughts to explain their observed tapeworm population differences in Alaska, when they saw more fine scale structuring than expected. Future field surveys on the genomics of different tapeworm life stages found at different environments and hosts should validate this premise.

Isolation by Distance (IBD) seems to be a driving factor in the tapeworm genetic differences among lakes (figure 5). Watershed also seems be a factor for genetic differences (figures 3 and 4) because it might be correlated to (or simply be a feature of) geographical distance. However, some lakes that belong to the same watershed but that are of some distance apart are also genetically different. For example, Lawson and Echo Lakes are further apart and genetically different to Gosling, Higgins, Boot, and Merrill lakes (table 3, figure 1), but all of them are from the same Campbell River Watershed. The R program ConStruct was not able to disentangle IBD from discrete models in describing the groupings of the data after K = 3 (figure 6). It is possible that both factors are contributing to the genetic structuring of the parasite. In fact, IBD, watershed, and host selection mentioned earlier, plus dispersal by birds might all contribute in some degree to the patterns of tapeworm genetic structuring seen in this work.

In summary, my hypothesis that the tapeworm should display little to no genetic structuring in Vancouver Island due to it having a highly mobile bird host (in accordance with the vagile host hypothesis) mostly holds true. The genetic differences between the populations were small; however, they were still significantly different from each other except for those populations that were close in distance or were from the same lake but collected in different years. And the most mobile bird host seems to have contributed mostly to the dispersal of the parasite and playing a major role on its genetic structuring seen here, but factors like less mobile hosts selection, IBD, and physical factors from each watershed might have also contributed, albeit in a lesser proportion, the genetic structuring seen in the tapeworm.

## Supplementary material

**Supplementary table 1:** SNPs with outlier F_ST_ obtained from Arlequin v3.5.2.2 based on 50,000 coalescent simulations (see Arlequin manual for more details).

| count | Locus | Obs.Het.BP | Obs.FST | FST.Pvalue | minus.log.Fst | 1FST.quantile |
| --- | --- | --- | --- | --- | --- | --- |
| 1 | 1624 | 0.621786 | 0.00699 | 1.06E-194 | 193.9735811 | 1 |
| 2 | 3058 | 0.610556 | 0.01841 | 4.51E-183 | 182.3455212 | 1 |
| 3 | 3242 | 0.627834 | 0.04102 | 4.11E-161 | 160.3864573 | 1 |
| 4 | 369 | 0.607571 | 0.03147 | 2.77E-106 | 105.5580238 | 1 |
| 5 | 4490 | 0.593525 | 0.00518 | 1.11E-73 | 72.95472397 | 1 |
| 6 | 3927 | 0.60401 | 0.0849 | 1.88E-69 | 68.72567817 | 1 |
| 7 | 1569 | 0.584581 | 0.00152 | 1.83E-61 | 60.73846117 | 1 |
| 8 | 8006 | 0.584489 | 0.02512 | 7.64E-49 | 48.11663671 | 1 |
| 9 | 7957 | 0.580216 | 0.01769 | 9.87E-40 | 39.00561817 | 1 |
| 10 | 4187 | 0.266214 | 0.62546 | 1.29E-34 | 33.89019542 | 1.29E-34 |
| 11 | 6885 | 0.578422 | 0.02662 | 2.31E-32 | 31.63659676 | 1 |
| 12 | 4825 | 0.614904 | 0.23605 | 1.58E-30 | 29.80263673 | 1 |
| 13 | 5159 | 0.520294 | 0.46245 | 1.94E-16 | 15.71231022 | 1.94E-16 |
| 14 | 6290 | 0.569792 | 0.0405 | 7.60E-16 | 15.11931729 | 1 |
| 15 | 2502 | 0.56627 | 0.03102 | 2.85E-14 | 13.54578342 | 1 |
| 16 | 1362 | 0.567377 | 0.03565 | 4.29E-14 | 13.36776143 | 1 |
| 17 | 4712 | 0.578223 | 0.08702 | 2.13E-13 | 12.67169584 | 1 |
| 18 | 4797 | 0.217377 | 0.53536 | 6.42E-10 | 9.192159315 | 6.42E-10 |
| 19 | 4103 | 0.554013 | 0.02759 | 1.04E-05 | 4.983392812 | 0.99999 |
| 20 | 3497 | 0.206141 | 0.40937 | 1.67E-05 | 4.777965412 | 1.67E-05 |
| 21 | 5595 | 0.55029 | 0.01723 | 3.96E-05 | 4.402511043 | 0.99996 |

**Supplementary table 2:** DAPC alpha-scores for different K values and their percent variance explained from the dataset. Alpha-scores measures the optimal number of Principal Components (PC) for each K value without overfitting the discriminant functions.

| K value | Suggested optimal number of PCs<sup>2</sup> | Percent variance explained<sup>3</sup> |
| --- | --- | --- |
| 2 | 1 | 3.6% |
| 3 | 24 | 18.7% |
| 4 | 25 | 19.2% |
| 5 | 25 | 19.2% |

<sup>1</sup>Suggested by Adegenet v2.1.5 to avoid overfitting the discriminant functions. PC stands for Principal Components.
<sup>2</sup>Genetic variance explained by the suggested optimal number of PCs from previous column

**Supplementary figure 1:**
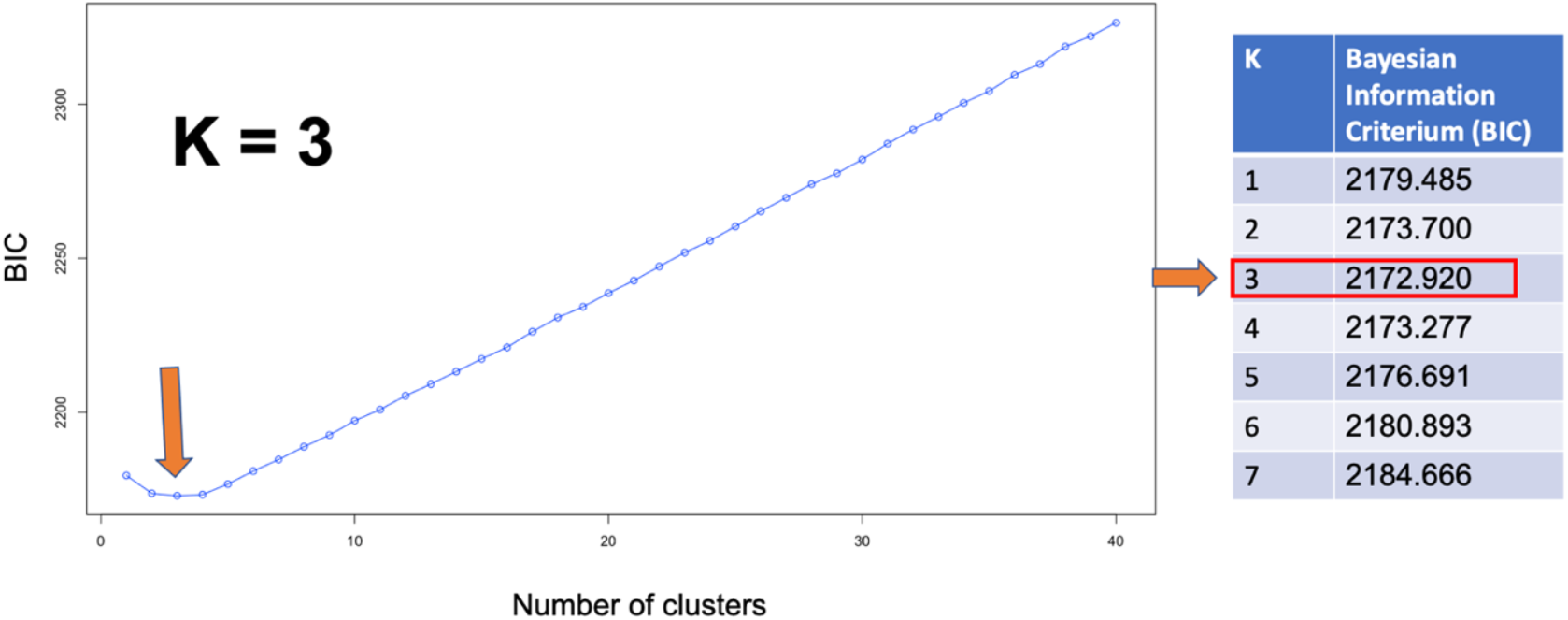
Best K value of 3 (indicated by orange arrow in graph) suggested by DAPC (from the Adegenet program). The lowest BIC number indicates the best K value for my dataset, and this number is indicated on the right-side table.

## #Comparing slopes between stickleback and tapeworm F_ST_

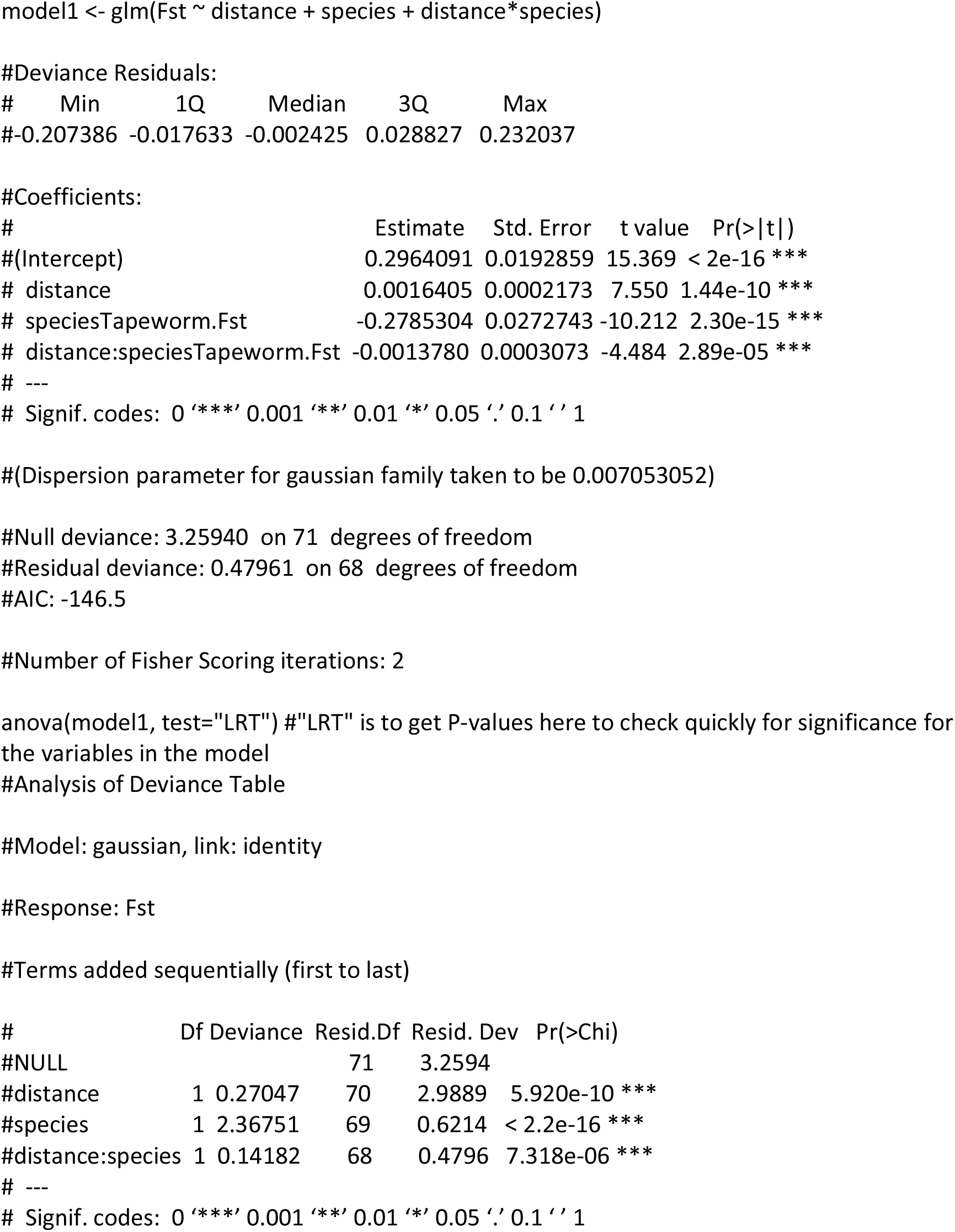

